# Rapid remodeling of the human gut microbiome in response to short-term animal product restriction

**DOI:** 10.1101/2025.07.30.667695

**Authors:** Christina Emmanouil, Maria Anezaki, Alexandros Simistiras, Stavros Glentis, Nikolaos Scarmeas, Pantelis Hatzis, Konstantinos Rouskas, Antigone S. Dimas

## Abstract

Diet strongly influences the gut microbiome, which in turn influences health, yet the effects of dietary patterns on microbiome composition and function remain underexplored in humans. We profiled a unique group of apparently healthy individuals from Greece, who alternate between omnivory and restriction of animal products for religious reasons (periodically restricted group, N=200). Using 16S rRNA sequencing, plasma metabolomics, and proteomics, we assessed the impact of three-to-four weeks of dietary restriction on gut microbiome composition and function and also explored links with host plasma biology. We compared findings to a continuously omnivorous group profiled in parallel (non-restricted group, N=211). Animal product restriction was found to reduce microbial diversity, primarily affecting rare taxa, and altered the abundance of nearly one-third of bacterial genera. Functional shifts included downregulation of cholesterol biosynthesis and purine degradation pathways, alongside upregulation of microbial biosynthesis of vitamin B2 and tryptophan, suggesting compensatory microbial responses to dietary nutrient depletion. Multi-omic integration revealed four microbial-metabolite-protein clusters, including a diet-responsive module linking *Negativibacillus* with potent metabolic regulator FGF21 and intermediate-density lipoproteins. Our findings demonstrate rapid adaptive plasticity of the human gut microbiome in response to short-term dietary restriction and highlight candidate microbial and molecular pathways that may mediate effects of animal product restriction on health.

## Introduction

The co-evolution of mammals and their microbial communities has given rise to complex symbiotic relationships defined by the bidirectional exchange of resources essential for host development, metabolism, and immune regulation^1^. In humans, the gut microbiome plays a central role in maintaining health, contributes to disease development^2,3^, and is strongly shaped by dietary intake^4,5^. Through the fermentation of proteins and indigestible plant polysaccharides, gut microbes produce a broad spectrum of metabolites, including essential vitamins, amino acids, and short-chain fatty acids (SCFAs), that influence inflammation, immune responses, energy homeostasis, and neurotransmission^1,6^. Disruptions in microbial composition and function have been associated with a range of diseases, including obesity, type 2 diabetes, cardiovascular disease, neurodegenerative disorders, and cancer^2,3,7,8^.

Dietary interventions are increasingly recognized as effective strategies for the prevention and management of chronic diseases, in part through their impact on the gut microbiome. However, investigating the effects of diet on the microbiome and human health remains challenging due to the complexity of dietary behaviors and interindividual variability^9^. Diets rich in animal products, particularly red meat, are characterized by increased protein fermentation and enrichment of bile-metabolizing bacteria such as *Alistipes putredinis*, *Bilophila wadsworthia* and *Ruminococcus torques*^4,10^. These compositional shifts are often linked to the production of inflammatory metabolites, including trimethylamine N-oxide (TMAO), and have been implicated in adverse cardiometabolic outcomes^8,11^.

Plant-based diets, typically rich in fiber, polyphenols, and complex carbohydrates, have been associated with reduced oxidative stress, reduced levels of low-grade inflammation, and improved metabolic health^12^. These dietary patterns are known to promote the enrichment of polysaccharide-fermenting and SCFA-producing gut bacteria, such as *Butyricicoccus* sp. and *Roseburia hominis*^10,12^, microbial shifts generally linked to favorable health outcomes, such as reduced inflammation, lower risk of obesity, type 2 diabetes, cardiovascular disease, and certain cancers^5,13,14^. However, plant-based dietary patterns have also been associated with lower intake of certain micronutrients, particularly vitamins B2, B12, and D, as well as calcium, and with potential health risks such as increased stroke incidence and lower bone mineral density^13,14^.

In the present study we performed 16S gut microbiome profiling of a unique group of apparently healthy individuals who voluntarily alternate between periods of omnivory and dietary restriction of animal products for religious reasons. The structured and predictable nature of this dietary pattern offers a naturalistic model that closely approximates a controlled dietary intervention. Microbiome profiles were compared between dietary states, and to profiles obtained in parallel from a control group of continuously omnivorous individuals. We uncovered dynamic responses of the microbiome to short-term animal product restriction, marked by a reduction in microbial diversity and by shifts in microbial abundance and metabolic activity. Furthermore, our analyses provide insights into the metabolic crosstalk between the gut microbiome and host plasma biology. Collectively, these findings enhance our understanding of human microbiome plasticity in response to dietary intake, highlight links between microbes and host biological pathways relevant for health, and may inform future development of microbiome-targeted therapeutic strategies.

## Methods

### Population sample and study design

A detailed description of the FastBio (religious <u>Fast</u>ing <u>Bio</u>logy) population sample has been outlined previously^15–17^. Briefly, participants who met selection criteria (**Supplementary Text 1**) belonged to one of two groups, specified by their diet. Periodically restricted individuals (PR, N=200, age: 20-76 years, sex: 54% female, BMI: 28.4±4.6 Kg m^-2^, **Supplementary Table 1**) alternate between omnivory and abstinence from meat, fish, dairy products and eggs for a total of 180-200 days annually, for religious reasons. Abstinence is practiced during four extended periods throughout the year, as well as on Wednesdays and Fridays of each week (**Fig. 1A**). Non-restricted individuals (NR, N=211, age: 19-74 years, sex: 55% female, BMI: 26.2±4.4 Kg m^-2^**, Supplementary Table 1**) are a continuously omnivorous, control group. Participants were invited to two scheduled appointments at the InterBalkan Hospital of Thessaloniki. The first appointment (T1 in autumn) was during a period of omnivory for both dietary groups. The second (T2 in spring) took place after three-to-four weeks of dietary restriction for the PR group, during Lent (**Fig. 1A and B**). Typically during this period, in addition to a pronounced reduction of animal fat intake, PR individuals undergo restriction of protein intake (as a proportion of total energy intake)^18,19^. This is driven chiefly by abstinence from all sources of animal protein and is not accompanied by changes in total energy or fiber intake. Our study design defines four dietary group by time point combinations or contexts (PR at T1, PR at T2, NR at T1, NR at T2).

**Figure 1.**
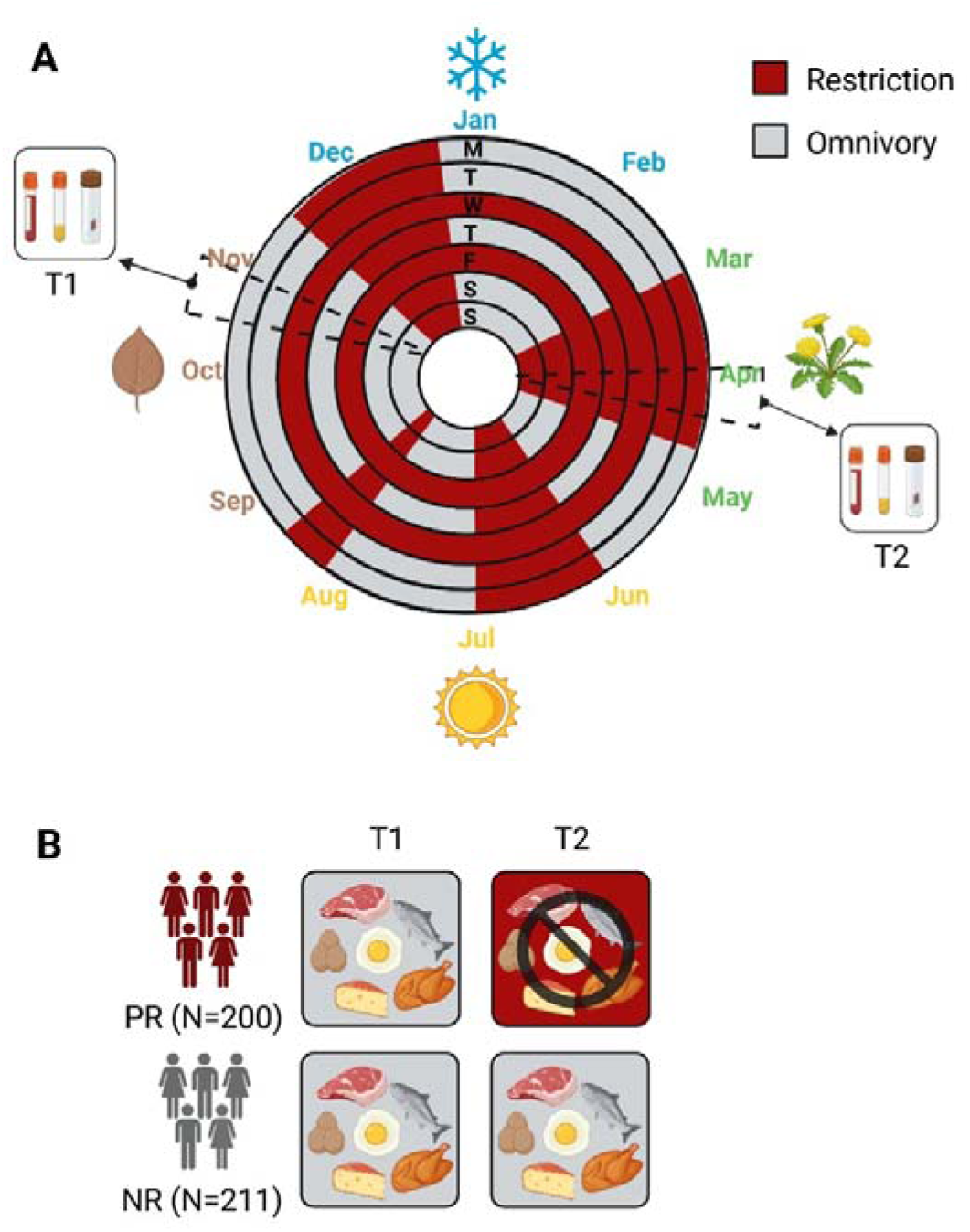
Periods of animal product restriction and study design. **A.** PR individuals practice animal product restriction for 180-200 days annually. Restriction is practiced during four extended periods throughout the year and on Wednesdays and Fridays of each week. **B.** FastBio study design. Periodically restricted (PR) individuals alternate between omnivory and animal product restriction for religious reasons. Participants were profiled at T1 (in autumn) and T2 (in spring). Continuously omnivorous individuals (NR group) were profiled in parallel, as a control group. Stool samples were delivered by 199 of the 200 PR participants and by all 211 NR participants. Created with BioRender.com

### Collection of biological material and measurement of traits

Participants collected two stool samples at home up to 48 hours prior to their appointment for both sampling time points. Samples were temporarily stored at - 20°C and were delivered by participants to the Interbalkan Hospital in an ice pack on the day of their appointment, where material was immediately stored at -80°C. Participants also submitted a completed Bristol score card linked to their sample. Individuals who had taken antibiotic, antifungal, antiviral or antiparasitic medication six months prior to T1 were excluded (**Supplementary Text 1**), while participants who had taken this type of medication between T1 and T2 were not excluded from the study, but information on medication use was recorded. The total of FastBio participants were also profiled for 16 blood biomarkers^16^, and for 249 and 1,455 plasma metabolites and proteins respectively (Nightingale Health Plc, Olink Explore 1536)^15^. Stool samples were delivered by 199 out of 200 PR and all 211 NR individuals. All participants provided written informed consent and the study was approved by the local ethics committee (BSRC Alexander Fleming Bioethics Committee Approval 17/02/2017).

### DNA extraction, amplification and sequencing

Microbial DNA was extracted from fecal samples using the QIAamp PowerFecal Pro DNA Kit (QIAGEN), following the manufacturer’s protocol. Briefly, 200 mg of fecal material was transferred to a 2 mL bead-beating tube, and lysis buffer was added. Samples were homogenized for 20 minutes using a Vortex-Genie 2 equipped with the QIA.13000-V1-24 adapter. For each extraction batch, a community standard sample of known composition (ZymoBIOMICS® Microbial Community Standard #D6300) was included as a control. DNA purity and concentration were quantified with Nano Drop and Qubit. Library preparation and 16S rRNA sequencing were performed at Oxford Genomics. The V3-V4 region of the 16S rRNA gene was targeted. Library preparation was performed as described in the Illumina workflow (#15044223 Rev.B). PCR amplicons were purified using Agencourt AMPure XP beads (Beckman Coulter, USA). Indexed amplicons were prepared (Nextera XT Index Kit), pooled in equimolar concentrations, denatured and diluted with a PhiX control and sequenced on a MiSeq system (2×300bp paired-end reads).

### Processing of 16S rRNA gene reads and taxonomic profiling

Computational analyses were performed in R v4.3.2^20^ and a detailed workflow can be found in **Supplementary** Fig. 1. Quality control on the sequencing reads was performed with FastQC^21^ and MultiQC^22^. Reads were trimmed with Cutadapt^23^ to remove sequencing adapters and chimeric PCR constructs. Reads mapping to the human genome (GRCh38) were removed using BWA^24^. Further processing was performed using the DADA2 algorithm^25^. Reads that matched the phiX genome were discarded. An ASV (Amplicon Sequence Variants) table was produced and taxonomy was assigned using the SILVA SSU Ref NR99 v138.1 database^26^, formatted for DADA2^27^. The minimum bootstrap confidence for assignment was set to 50. Assignment as *Incertae sedis* was replaced by NA. We retained only reads with assigned taxonomy at genus level, and samples with data from both time points (paired samples; 188 PR, 194 NR). Each sample yielded an average of 111,483 reads, and a total of 26,855 unique ASVs were retained for downstream analysis.

### Quantification of alpha and beta diversity

We rarefied the ASV table (function “rarefy_even_depth”) to control for differential sequencing outputs. The rarefaction threshold was 64,000 reads and was chosen to maximize sequencing depth while losing only four individuals (2 PR, 2 NR). Diversity measures were calculated using the R packages “phyloseq”^28^ and “vegan”^29^ (functions “estimateR”, “diversity”, “ordinate”, “adonis2”). Alpha-diversity was assessed through four indices which reflect the richness (Observed richness), balance (Pielou’s evenness) and diversity (Shannon and Gini-Simpson indices) of the bacterial community. We tested for differences in alpha-diversity between contexts using a Wilcoxon signed-rank test (across time points) or a Wilcoxon rank-sum test (between dietary groups) (p<0.05). For beta-diversity, the distance matrix was calculated using the Bray-Curtis dissimilarity index and plotted using principal coordinate analysis (PCoA). Variance explained by each explanatory variable was calculated with PERMANOVA^30^, using a sequential model and 999 permutations. The p-values were adjusted for multiple testing using the Benjamini-Hochberg method (q<0.05) and significant factors were used as covariates in subsequent analyses.

### Differential abundance analysis of genera and of predicted metabolic pathways

Using the unrarefied ASV table, we agglomerated ASVs to genus level (function “tax_glom”), filtered for abundance (0.0001%) and prevalence (10%) and rarefied to 60,000 reads. Filtering thresholds were the same for all analyses. To interrogate differences in bacterial abundance between contexts, we normalized abundances to the range [0,1] and employed a CPLM model in MaAsLin2^31^ with time point and selected covariates (**Supplementary Text 2**) as fixed effects, and participant ID as a random effect. The p-values were adjusted for multiple testing using the Benjamini-Hochberg method (q<0.05). For functional profiling of the bacterial community, we started with the unrarefied ASV table, performed abundance and prevalence filtering as described above, rarefied to 48,000 reads and employed PICRUSt2 to predict functional abundances^32^. Predicted MetaCyc pathways were subsequently filtered for abundance and prevalence. Changes in pathway expression between contexts were tested using MaAsLin2 as described above, using an LM model.

### Links between abundance of bacterial genera and of microbial pathways with clinical health markers

To investigate potential connections between gut microbiome shifts and health-related markers, we explored correlations between blood biomarker levels from^16^ and diet-associated changes in: a) bacterial abundance, and b) expression of predicted microbial pathways. We regressed out effects of covariates (**Supplementary Text 2**), calculated Spearman’s correlation coefficient (“stats” package, function “cor.test”) and corrected for multiple testing using the Benjamini-Hochberg method (q<0.05).

### Links between abundance of bacterial genera and molecular phenotypes

To explore links between microbiome composition and host plasma biology, we examined associations between bacterial abundance and plasma metabolite and protein levels. We added a pseudo-count of 1 to bacterial abundance and performed centered log-ratio transformation to mitigate effects of compositionality. Missing metabolite and protein values were imputed with mean value imputation^15^ and covariate effects were regressed out (**Supplementary Text 2**). DIABLO^33^ (“mixOmics” package^34^) was used, with a full-weighted design matrix. To account for repeated measures, we extracted the within-subject variance. The optimal number of components was defined using 50x5-fold cross-validation and the optimum number of variables per component per dataset was determined with 5x10-fold cross-validation. Clusters encompassing at least one of each biological levels (microbiome, metabolites, proteins) and displaying a correlation of r≥|0.6| were visualized.

## Results

### Population sample

We profiled a unique group of apparently healthy individuals from Greece who alternate between omnivory and dietary restriction of animal products as part of religious fasting, and compared findings to a control group of continuously omnivorous individuals profiled in parallel (**Fig. 1, Supplementary Table 1**). Participants were profiled at two time points: T1 in autumn, during a period of omnivory for both dietary groups, and T2 in early spring, following three-to-four weeks of animal product restriction for PR individuals, during Lent. Our study design therefore involves four dietary group by time point combinations (contexts). We have previously shown that this type of short-term dietary restriction induces broad immunometabolic reprogramming, with mostly positive effects on health, marked by reductions in plasma levels of cholesterol and other lipid classes, of branched-chain amino acids, and shifts in immunometabolic regulators, including a pronounced increase in potent metabolic regulator FGF21^15,16^. In the present study we explored the effects of this dietary pattern on the gut microbiome using 16S rRNA gene sequencing and combined with findings from plasma biomarkers, metabolites and proteins^15,16^.

### Taxonomic profiling and diet-responsive shifts in bacterial diversity

The gut microbiomes of participants were predominantly composed of bacteria from Lachnospiraceae, Ruminococcaceae, and Bacteroidaceae families, and *Bacteroides*, *Faecalibacterium*, and *Blautia* genera (quantified as average abundance per sample; **Supplementary** Fig. 2A). While the relative abundances of the 20 most prevalent genera were broadly similar across groups and time points (**Supplementary. Fig. 2B**), as expected we found substantial variability between individual participants (**Supplementary** Fig. 2C)^9^. Short-term animal product restriction was associated with a reduction in microbial diversity as measured by observed richness and the Shannon diversity index (Wilcoxon, p<0.05), while Pielou’s evenness and the Gini-Simpson index remained unchanged (Wilcoxon, p>0.05) (**Fig. 2**). These findings suggest that animal product restriction, which involves reduced dietary diversity, may primarily lead to loss of low-abundance taxa. In the control group, microbial diversity remained stable between T1 and T2 (Wilcoxon, p>0.05; **Fig. 2**).

**Figure 2.**
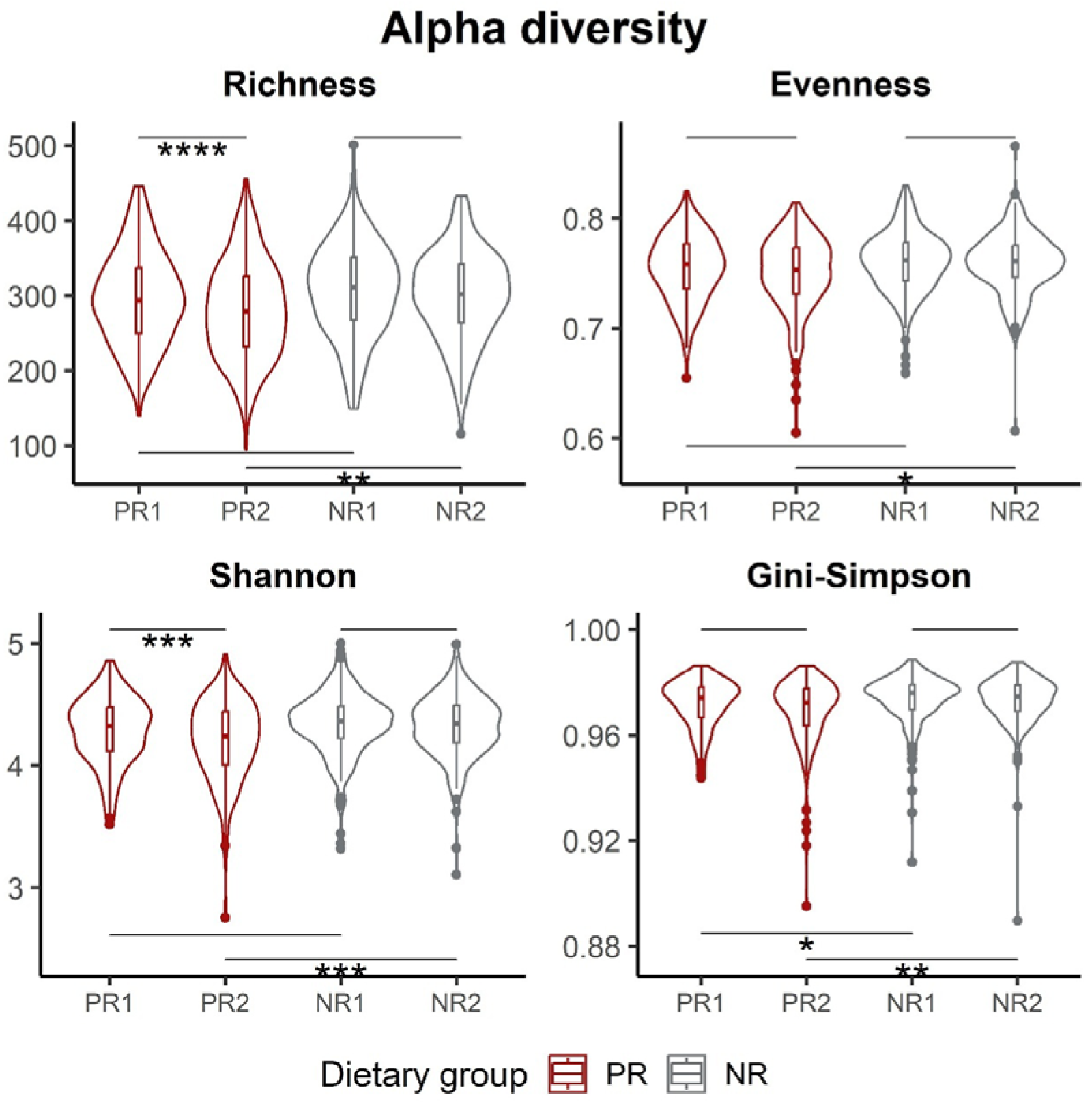
Alpha diversity metrics for each dietary group by time point combination (context). Violin plots and boxplots show the distribution of Observed richness, Pielou’s evenness, and Shannon and Gini-Simpson’s diversity indices. The lower and upper hinges of the boxplots correspond to the first and third quartiles. Statistical comparisons were performed using the Wilcoxon signed-rank test between paired samples (PR1-PR2, NR1-NR2) and the Wilcoxon rank-sum test between independent samples (PR1-NR1, PR2-NR2). Asterisks indicate levels of significance (*p<0.05, **p<0.01, ***p<0.001, ****p<0.0001). PR1: PR at T1, PR2: PR at T2, NR1: NR at T1, NR2: NR at T2.

We also explored differences between dietary groups and found that when both groups were omnivorous (T1), microbial diversity was largely comparable, with the exception of lower Gini-Simpson diversity in the PR group (Wilcoxon, p<0.05; **Fig. 2**), possibly reflecting subtle differences in a few dominant taxa that likely do not affect overall composition and balance. However, during dietary restriction (T2), the PR group exhibited substantially lower microbial diversity across all tested indices compared to the control group (Wilcoxon, p<0.05; **Fig. 2**). Microbial community structure also differed by dietary group and state, as determined by PERMANOVA on Bray-Curtis distances (R^2^=0.0063, q<0.05) and was additionally associated with age^2^, sex, BMI, Bristol score, medication use, and smoking (R^2^=0.0019-0.0058, q<0.05; **Supplementary** Fig. 3).

### Effects of animal product restriction on bacterial abundance and predicted metabolic pathways

Dietary restriction altered the relative abundance of nearly one-third of bacterial genera tested (47 out of 161), while in the control group only *Lactiplantibacillus* was differentially abundant at T2 (**Fig. 3A and B, Supplementary Table 2**). Three quarters of diet-associated changes involved a decrease in abundance, with the direction of change being largely consistent within taxonomic families (**Supplementary Table 3**). Notably, lactic acid bacteria (LAB) exhibited consistent reductions in abundance, driven chiefly by changes in Lactobacillaceae (*Lactiplantibacillus*, *Lacticaseibacillus*, *Lactobacillus*, *Latilactobacillus*), but also in *Enterococcus*, *Lactococcus*, *Streptococcus.* LAB are commonly derived from fermented dairy products and include strains with probiotic properties^35,36^. Changes were also found for SCFA-producing genera, but with variable direction (e.g. increased *Butyricicoccus* and *NK4A214 group*, but decreased *Holdemania* and *Blautia*). Overall, we found that the most pronounced shifts occurred in less abundant families, while dominant families such as Bacteroidaceae, Prevotellaceae, and Bifidobacteriaceae remained largely stable in response to dietary restriction.

**Figure 3.**
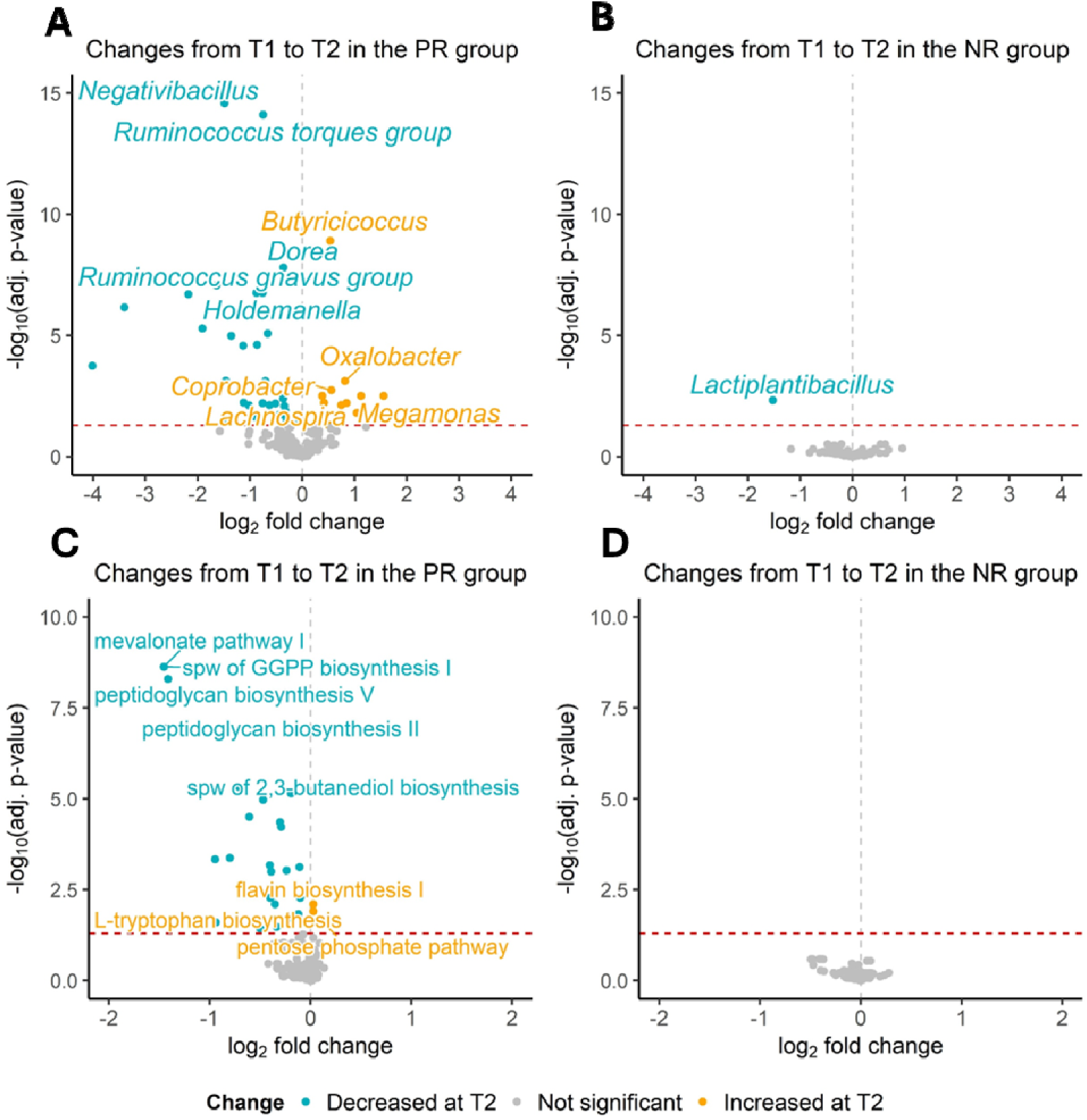
Differentially abundant genera and pathways across time points. Volcano plots displaying log_2_ fold changes and -log_10_ p-values (BH-adjusted) for bacterial genera between time points in the PR group (**A**) and in the NR group (**B**), and for microbially expressed predicted pathways between time points in the PR group (**C**) and in the NR group (**D**). Significant changes (p<0.05) are highlighted in blue (decreased at T2) or yellow (increased at T2) color. Spw: superpathway.

We next investigated the functional consequences of diet-responsive shifts in microbial composition and found that 9% (27 of 302) of predicted pathways were affected, with nearly all changes reflecting downregulation of microbial functions (**Fig. 3C, Supplementary Table 2**). No changes were found in the control group (**Fig. 3D, Supplementary Table 2**). The most prominently downregulated pathways were related to cholesterol biosynthesis (mevalonate pathway I, superpathway of GGPP biosynthesis I (via mevalonate)), a nutrient that is substantially depleted during dietary restriction of animal products. Downregulation was also found for pathways involved in bacterial cell wall synthesis (peptidoglycan biosynthesis V and II), likely reflecting a reduction in bacterial biomass in accordance with the overall decrease in microbial abundance. Additionally, purine degradation pathways were downregulated (purine nucleobases degradation I, guanosine nucleotides degradation III, purine nucleotides degradation II, adenosine nucleotides degradation II and IV), possibly reflecting a microbial response toward nitrogen conservation under animal product restriction. In contrast, a small number of pathways were upregulated, including flavin biosynthesis I and L-tryptophan biosynthesis, involved in the production of essential nutrients (vitamin B2 and tryptophan, respectively) that are primarily obtained from animal-derived foods^37,38^. The pentose phosphate pathway (PPP), which generates NADPH and ribose-5-phosphate for anabolic metabolism and nucleotide synthesis^39^, was also upregulated following short-term restriction.

When comparing compositional and functional profiles between dietary groups, we found that profiles were identical when both groups consumed an omnivorous diet, with the exception of slightly higher abundance of *Faecalitalea* in the PR group. However, three-to-four weeks of animal product restriction resulted in divergence of microbiomes between dietary groups, with differences found for 14 genera and 12 metabolic pathways (**Supplementary** Fig. 4A, B, C **and D, Supplementary Table 2**). The majority of these differences reflected lower abundance or downregulation in the PR group. Notably, over half of these differences were also detected in the PR group from T1 to T2, suggesting direct effects of dietary restriction on microbial composition and function.

### Associations of bacterial abundance and of metabolic pathways with clinical health markers

We next examined associations between genera influenced by dietary restriction and clinical health markers (**Fig. 4A**). A pattern spanning a wide range of bacterial taxonomic groups, was found for correlations of changes in bacterial abundance with levels of triglycerides, insulin, urea, ALT and CRP. The genera underlying most of the correlations were *Ruminococcus gnavus group* (Lachnospiraceae), *NK4A214 group* (Oscillospiraceae) and *Megamonas* (Selenomonadaceae). Similarly, changes in predicted microbial pathways correlated with health markers, including levels of glucose, insulin, urea, creatinine, ALT, and gGT (**Fig. 4B**). Notably, half of these associations were found for urea, which was mostly linked to carbohydrate or amino acid metabolism pathways.

**Figure 4.**
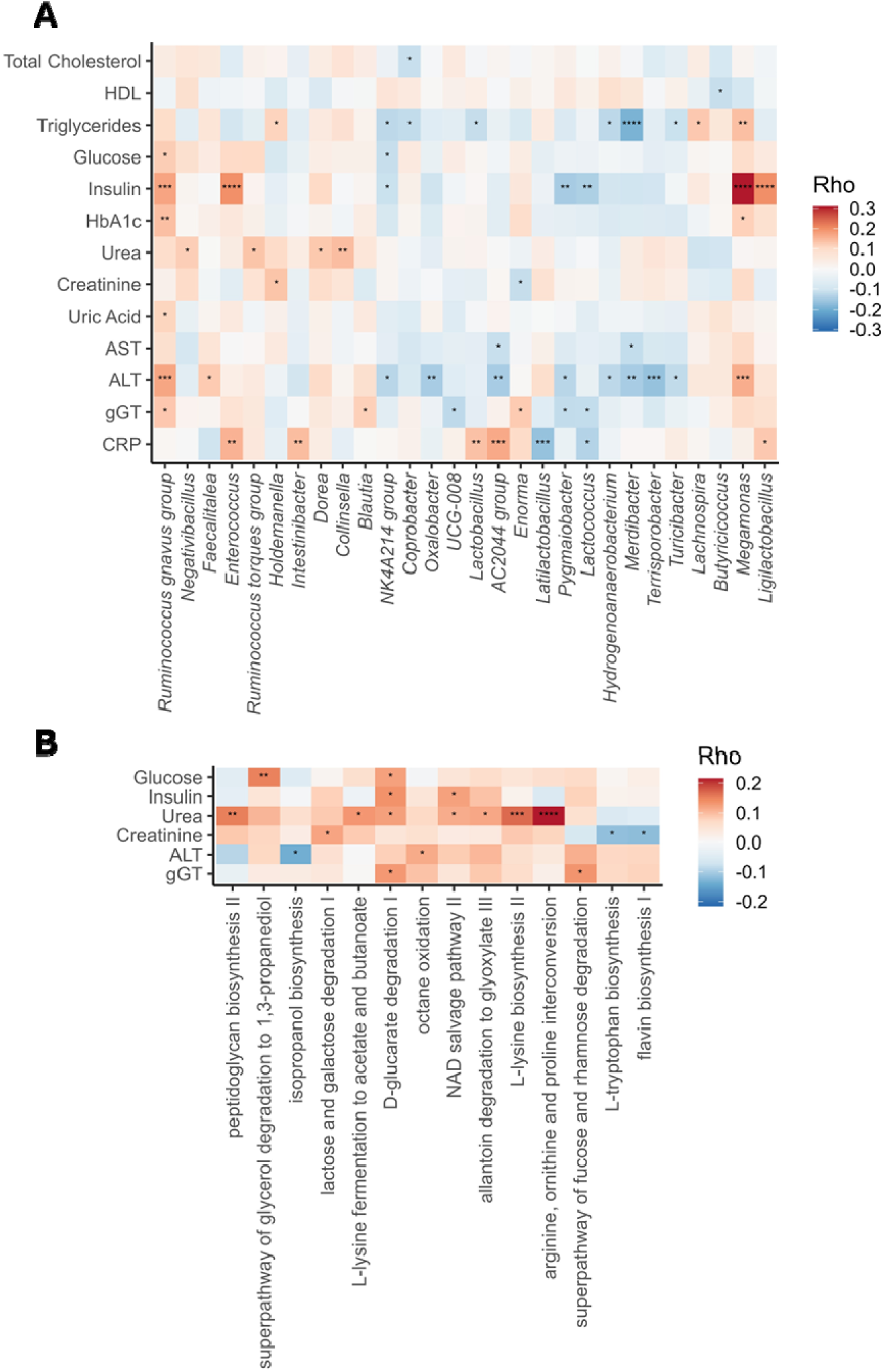
Correlations of bacterial genera with clinical markers of health. Heatmaps showing Spearman’s correlations of levels of clinical health markers with bacterial abundance (**A**) and with expression of predicted metabolic pathways (**B**). Only diet-responsive, differentially abundant genera and pathways were included in the analysis. We display markers, genera and pathways with at least one significant correlation. P-values were BH-adjusted and asterisks indicate levels of significance (*p<0.05, **p<0.01, ***p<0.001, ****p<0.0001).

### Multi-omic integration of microbiome, metabolomic, and proteomic data

To investigate potential connections between gut microbiome composition and plasma biology, we integrated microbiome, metabolomic and proteomic data and identified four bacterial-protein-metabolite clusters (**Fig. 5A and B**). The first cluster was diet-responsive, consisting of entities that independently exhibited marked changes in response to animal product restriction. *Negativibacillus*, the genus showing the most substantial diet-responsive decrease in abundance, was associated with FGF21, a protein that increased 2.4-fold in abundance, as well as to HAVCR1 and SPON2, both among the most diet-responsive proteins, and to IDL and XS VLDL, lipoprotein subclasses which were markedly decreased during dietary restriction^15^.

**Figure 5.**
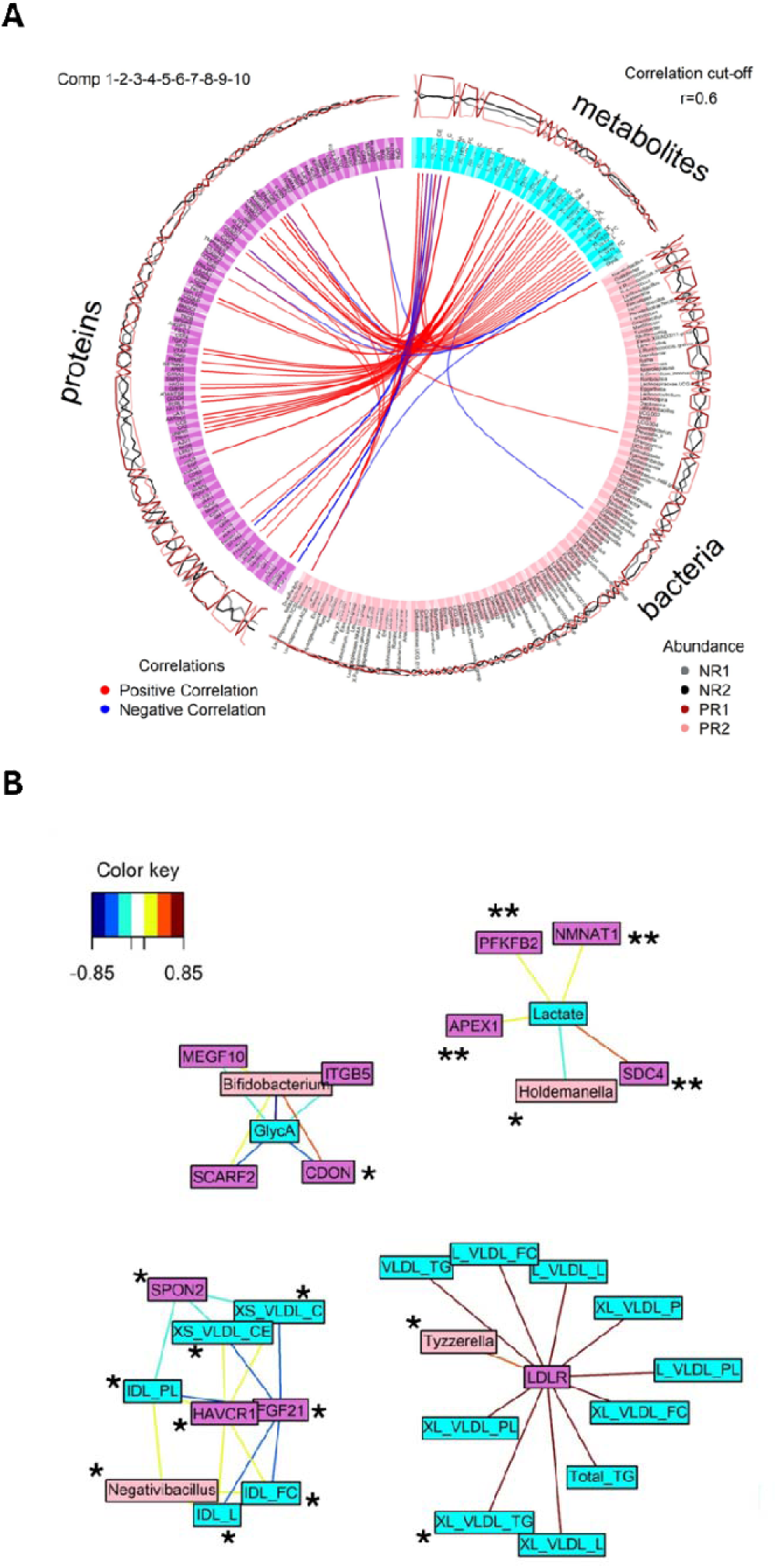
Correlations between bacterial genera and plasma metabolites and proteins. **A.** Circos plot indicating the connectivity between all three datasets (|r|>0.6). Blue lines indicate negative correlations, red lines indicate positive correlations. The four lines at the outer side of the circle indicate the abundance of each bacterium or molecule in each of the four contexts. PR1: PR at T1, PR2: PR at T2, NR1: NR at T1, NR2: NR at T2. **B.** Network indicating correlations between bacteria, metabolites and proteins. Light pink nodes represent genera, blue represent metabolites and dark pink represent proteins. Single asterisks indicate entities with changes in abundance in the PR group from T1 to T2. Double asterisks indicate entities with changes in abundance in both the PR and the NR group from T1 to T2. Clusters encompassing at least one of each biological levels (microbiome, metabolites, proteins) and displaying a correlation of r≥|0.6| were visualized. Correlation values are indicated with a color spectrum from blue (-0.85) to red (0.85).

The second and third clusters included bacterial genera influenced by diet and associated with proteins and metabolites showing mixed responses. *Holdemanella* was linked to lactate, and to proteins APEX1, NMNAT1, SDC4, PFKFB2, which are primarily involved in glycolysis and redox regulation. *Tyzzerella* was associated with LDLR, and with L VLDL and XL VLDL particles, indicating a potential link with cholesterol metabolism. Components of the fourth cluster did not display diet-responsive effects and included *Bifidobacterium*, which was linked to GlycA, SCARF2, CDON, ITGB5 and MEGF10, molecules primarily related to inflammatory processes.

## Discussion

In the present study we performed gut microbiome profiling in a unique group of apparently healthy individuals who follow a consistent and predictable pattern of alternating between omnivory and restriction of animal products for approximately 180–200 days annually. To our knowledge, this is the first study to explore microbiome plasticity in response to a periodic, real-world dietary restriction pattern sustained over more than a decade, while also integrating with host plasma biology, an often-underexplored dimension in microbiome research. We demonstrated rapid compositional and functional adaptation of the microbiome in response to animal product restriction, and identified correlations between bacterial genera, clinical health markers, and molecular phenotypes.

We found that short-term restriction of animal products led to a rapid reduction in microbial diversity, without affecting community evenness, likely driven by the selective loss of rare taxa. Previous studies examining microbial diversity under plant-based diets have reported mixed results^40^. However, more recent research in larger and better-characterized populations has demonstrated reduced diversity in individuals adhering to plant-based diets compared to omnivores^10,41,42^. Although reduced diversity is often observed in diseases of affluence, possibly due to the disruption of complex trophic networks^43^, we propose that in healthy individuals reduced diversity may reflect an adaptive response toward functional optimization. During short-term restriction of animal products, when a less diverse diet is being consumed, the microbiome may shift to align with altered nutrient availability, reflecting functional plasticity through loss of mostly rare taxa. Rare taxa have been proposed to constitute a reservoir of genetic potential, capable of supporting essential microbial functions under diverse conditions^44^. Therefore, their depletion under a less diverse diet may reflect an adaptive response that may not necessarily have negative consequences for health.

Although mostly rare taxa were lost following animal product restriction, we found that the abundance of nearly one-third of bacterial genera tested was altered, with most changes involving decreased bacterial levels. Genera typically associated with meat and dairy consumption including *Ruminococcus torques group*, *Ruminococcus gnavus group* and *Collinsella*, decreased, while genera commonly linked to plant-based diets, including *Lachnospira* and *Butyricicoccus*, increased^10,45–47^. However, certain fiber-fermenting genera frequently enriched in vegan diets, such as *Bifidobacterium*, *Anaerostipes* and *Roseburia*^10,12,48^, remained unaffected. It is plausible that this is because animal product restriction during Lent is typically not accompanied by changes in dietary intake of fiber. Nevertheless, changes in other fiber-fermenting genera were found, suggesting that the dynamics of these genera may be shaped not only by fiber availability, but also by factors such as microbial competition and ecological niche adaptation^49^.

A marked reduction was found in genera belonging to the LAB group, likely driven by the exclusion of fermented dairy products, the primary source of these bacteria^35^, during restriction. Indeed, certain LAB taxa, including *Streptococcus*, *Lactobacillus*, *Lacticaseibacillus*, *Lactococcus*, have been shown to be more abundant in omnivorous and vegetarian diets compared to vegan diets and are consistently associated with dairy intake^10,50,51^, although *Lacticaseibacillus* and *Streptococcus* have also been found to be enriched in individuals following high-protein diets^12^. As many LAB strains are considered probiotic due to their capacity to produce antioxidant, anti-inflammatory and immunomodulatory metabolites, their decrease may reflect a diet-induced shift that could influence host biology^52,53^.

We identified additional diet-associated shifts in bacterial abundance, including decreases in genera previously linked with adverse health outcomes. *Negativibacillus*, which was markedly reduced under dietary restriction, has been associated with inflammatory bowel disease (IBD)^54^, non-alcoholic fatty liver disease^55^ and treatment resistance in Crohn’s disease^56^. *Merdibacter*, associated with chronic kidney disease (CKD)^57^, IBD^58^ and irritable bowel syndrome^59^, and *Terrisporobacter*, associated with dyslipidemia^60^, also showed reduced abundance. However, some diet-associated shifts may reflect negative effects, such as increased abundance of *Megamonas*, a genus associated with metabolic syndrome^61^ and CKD^62^. To date, considerable inconsistency remains regarding associations between specific bacterial genera and human health, and it is often unclear whether changes in microbial abundance are a cause or consequence of disease. This underscores the challenge of defining a universal ‘healthy microbiome’ signature^63^. Nevertheless, the diet-responsive changes uncovered in the present study mostly involved reductions in genera previously associated with adverse health outcomes.

Overall, we found two broad types of microbiome metabolic responses to dietary change, which we refer to as passive and compensatory. The most pronounced passive response was the downregulation of the mevalonate pathway, which plays a key role in the synthesis of isoprenoids and sterols, including cholesterol. During animal product restriction, dietary intake of cholesterol is substantially reduced, a shift also reflected in decreased plasma levels of cholesterol and other lipid classes^15^. In the host, such a reduction typically leads to a compensatory response involving upregulation of the mevalonate pathway and of GGPP biosynthesis, to increase endogenous synthesis of cholesterol and cellular uptake^64,65^. Consistent with this, our previous work showed that mevalonate kinase, a key enzyme in this pathway^66^, increases in plasma abundance following dietary restriction^15^. In the gut microbiome, the mevalonate pathway is primarily expressed by gram-positive genera, such as *Enterococcus*, *Lactobacillus*, *Streptococcus* (all LAB taxa, that decreased during restriction) as well as *Staphylococcus*, while most gram-negative bacteria rely on the alternative methylerythritol phosphate (MEP) pathway to synthesize isoprenoids^67^. Thus, it is plausible that this reflects a passive response likely stemming from a nutrient-driven reduction of genera which employ this pathway.

We also identified compensatory responses of the microbiome to replenish nutrients depleted during animal product restriction. Most notably, we found upregulation of microbial biosynthetic pathways for tryptophan and vitamin B2 (riboflavin), both of which are essential nutrients primarily obtained from animal-derived foods^37,38^. Tryptophan serves as a precursor for key molecules such as serotonin, melatonin and niacin (vitamin B3), and plays a central role in the gut-brain axis, immune regulation, and energy metabolism^37^. Similarly, vitamin B2 is essential for energy metabolism, fatty acid oxidation, purine catabolism, redox balance, and for the metabolism of other B vitamins^38^. Additionally, we detected downregulation of purine degradation, and upregulation of the PPP, which produces ribulose-5-phosphate. This shift may also be linked to the upregulation of vitamin B2 biosynthesis, as its production requires both guanosine triphosphate (GTP) and ribulose-5-phosphate. These changes suggest a microbial response to sustain key metabolic functions and nutrient availability under dietary restriction.

When exploring links between the microbiome and health-related biomarkers, we identified correlations between diet-responsive genera and clinical markers, including triglycerides, insulin, urea, ALT, and CRP. Although diet-driven microbiome shifts were associated with changes in biomarkers associated with both positive and negative effects on health, we found a consistent link with improved markers of renal function, primarily through decreased levels of urea. This aligns with lower intake of dietary protein during the restriction period^68^ and may be reflected in the downregulation of purine degradation pathways during animal product restriction, suggesting a potential microbial response toward nitrogen conservation.

We next explored links between the gut microbiome and plasma biology and uncovered previously unreported associations between bacterial genera, plasma metabolites and proteins, identifying four distinct clusters. The first cluster emerged as a diet-responsive module comprising components that showed the most pronounced changes during dietary restriction. *Negativibacillus*, the genus with the greatest diet-associated decrease, is a recently discovered taxon^69^ whose elevated levels have been associated with resistance to Crohn’s disease treatment^56^, IBD^54^, and non-alcoholic fatty liver disease^55^.

The *Negativibacillus* cluster included associations with IDL and XS VLDL lipoproteins, as well as with FGF21, HAVCR1 and SPON2 proteins^15^. VLDL particles, secreted by the liver, transport triglycerides and cholesterol and are progressively converted into IDL through triglyceride depletion^70^. The XS VLDL subclass, having lost much of its triglyceride content, is similar in size and density to IDL, suggesting that this cluster may also capture effects related to lipoprotein particle size. Both VLDL and IDL are components of remnant cholesterol, an atherogenic lipid fraction associated with increased risk of CVD, including coronary heart disease, stroke, cardiac death, and all-cause mortality^71^. Of the protein components of this cluster, liver-secreted FGF21 is a hormone with roles in energy homeostasis, lipid and glucose metabolism and insulin sensitivity, and has been shown to extend lifespan in model organisms^72^. In mice, dietary protein restriction induces hepatic FGF21 expression, elevating circulating FGF21 levels in a response shown to be mediated by the gut microbiome^73^. HACVR1 is involved in regulation of immune cell activity and renal regeneration^74^, while SPON2 is involved in macrophage activation and has been associated with tumor progression^75^, whereas in mice it has been associated with immune response, inflammation and hepatic lipid metabolism^76^. Overall, this cluster appears to capture metabolic adaptations to animal product restriction, and highlights previously unreported connections for further investigation of their likely effects on immunometabolic and cardiometabolic homeostasis.

The other clusters comprised components with mixed responses to dietary restriction, suggesting interactions that are either unresponsive to this dietary shift, or vary in responsiveness across molecular levels. In the second cluster, *Holdemanella* was linked to lactate, to NMNAT1 and APEX1, proteins involved in NAD+ biosynthesis and the oxidative stress response^77,78^, to PFKFB2, which mediates lactate-dependent macrophage glycolysis during efferocytosis in mice^79^, and to SDC4, which has shown correlations with markers of oxidative stress and inflammation^80^. This cluster likely captures effects of redox regulation, potentially reflecting changes that underlie reduced oxidative stress typical of plant-based diets^12^. The third cluster included *Tyzzerella*, VLDL particles and LDLR, pointing to a potential link to cholesterol metabolism and cardiovascular health. This is consistent with findings linking *Tyzzerella* to CVD risk^81^ and Crohn’s disease^82^ and with the known role of LDLR in binding VLDL via apoB and apoE^83^.

The fourth cluster linked *Bifidobacterium* to GlycA, a marker of systemic inflammation, relevant to CVD risk (Ballout and Remaley, 2020), to CDON and SCARF2, proteins with likely associations to cardiovascular health^84,85^, and to ITGB5, a potential target in treating diabetic cardiovascular complications^86^. Indeed, Bifidobacteria have been shown to reduce levels of TMAO^87^, a metabolite typically elevated in meat-based diets and associated with CVD pathogenesis^8,11,88^. While MEGF10, also found in this cluster, has not been directly linked to cardiovascular health, it is a protein likely involved in inflammatory processes through its role in apoptotic cell clearance in the mammalian brain^89^. Although the components of this cluster did not show diet-responsive effects in their independent analyses, we propose that these links may reflect systemic effects of animal protein restriction that could potentially be captured in a larger study group.

Our study has several limitations. First, although the design captures the overall impact of animal product restriction, it does not enable us to disentangle effects stemming from specific dietary components, and therefore reflects changes associated with the absence of a broader set of nutrients rather than individual dietary factors. However, this limitation is offset by the value of studying real-world dietary patterns, which provides more insights than studying individual nutrients in isolation from the broader dietary context^90^. Second, the use of 16S rRNA gene sequencing limits taxonomic resolution to the genus level, and limits functional interpretation. Given the prominent effects of animal product restriction on gut microbiome composition and function we were nevertheless able to glean broader genus-level insights. Third, although we have highlighted links between bacteria and blood biomarkers, metabolites and proteins, these represent correlations and further work is required to establish functional connections and determine causal mechanisms. Finally, the metabolomic and proteomic platforms employed covered only a subset of plasma molecules, restricting our analyses to the molecules included in these assays.

In conclusion, we have demonstrated that short-term dietary restriction of animal products induces dynamic remodeling of the gut microbiome. Given the long-term adherence of participants to this dietary pattern, and considering that both dietary groups exhibited similar diversity, compositional, and functional profiles under omnivorous conditions, the response to dietary restriction appears to be rapid, but it is also most likely transient. While direct causal links between specific genera and health remain challenging to establish, we have highlighted diet-responsive taxa, including *Negativibacillus*, *Merdibacter*, and *Megamonas*, whose effects remain poorly understood, and suggest that they may serve as candidates for further functional investigation or for biomarker development. Additionally, we revealed correlations between microbial taxa and plasma metabolites and proteins, which can aid in hypothesis generation for causality testing. Overall, our findings suggest potential diet-responsive effects on health mediated through the gut microbiome and highlight candidate mechanisms of host-microbiome crosstalk. A deeper understanding of how the gut microbiome systemically adapts to dietary changes will shed light on the mechanisms by which diet shapes human health. These insights could inform future strategies for microbiome-targeted interventions and the development of pharmacological approaches that mimic the beneficial effects of dietary restriction.

## CRediT authorship contribution statement

Conceptualization: CE, ASD. Methodology: CE, AS, KR, ASD. Formal analysis: CE, KR. Investigation: CE, MA, SG, KR, ASD. Data curation/software: CE. Visualization: CE. Resources: ASD. Supervision: NS, ASD. Writing-original draft: CE, ASD. Writing-review and editing: CE, NS, PH, KR, ASD. Project administration: ASD. Funding acquisition: ASD

## Acknowledgements

The authors are grateful to the FastBio study participants and to the Interbalkan Hospital Staff. We would also like to thank Dr Dimitrios Rouskas, Dr Pavlos Rouskas and Dr Loukas Kipouros for their invaluable help with sample collection. We acknowledge the support of Dr Theodosios Kyriakou at Oxford Genomics, and of the Genomics Facility at BSRC Alexander Fleming. We would also like to thank Professor Mary Yannakoulia for useful discussions.

## References

1 Lynch, S. V. & Pedersen, O. The Human Intestinal Microbiome in Health and Disease. N Engl J Med 375, 2369–2379, doi:10.1056/NEJMra1600266 (2016).

2 Singh, R. K. et al. Influence of diet on the gut microbiome and implications for human health. J Transl Med 15, 73, doi:10.1186/s12967-017-1175-y (2017).

3 de Vos, W. M., Tilg, H., Van Hul, M. & Cani, P. D. Gut microbiome and health: mechanistic insights. Gut 71, 1020–1032, doi:10.1136/gutjnl-2021-326789 (2022).

4 David, L. A. et al. Diet rapidly and reproducibly alters the human gut microbiome. Nature 505, 559–563, doi:10.1038/nature12820 (2014).

5 Wilson, A. S. et al. Diet and the Human Gut Microbiome: An International Review. Dig Dis Sci 65, 723–740, doi:10.1007/s10620-020-06112-w (2020).

6 Canfora, E. E., Jocken, J. W. & Blaak, E. E. Short-chain fatty acids in control of body weight and insulin sensitivity. Nat Rev Endocrinol 11, 577–591, doi:10.1038/nrendo.2015.128 (2015).

7 O’Keefe, S. J. Diet, microorganisms and their metabolites, and colon cancer. Nat Rev Gastroenterol Hepatol 13, 691–706, doi:10.1038/nrgastro.2016.165 (2016).

8 Fan, Y. & Pedersen, O. Gut microbiota in human metabolic health and disease. Nat Rev Microbiol 19, 55–71, doi:10.1038/s41579-020-0433-9 (2021).

9 Lozupone, C. A., Stombaugh, J. I., Gordon, J. I., Jansson, J. K. & Knight, R. Diversity, stability and resilience of the human gut microbiota. Nature 489, 220–230, doi:10.1038/nature11550 (2012).

10 Fackelmann, G. et al. Gut microbiome signatures of vegan, vegetarian and omnivore diets and associated health outcomes across 21,561 individuals. Nat Microbiol, doi:10.1038/s41564-024-01870-z (2025).

11 Senthong, V. et al. Intestinal Microbiota-Generated Metabolite Trimethylamine-N-Oxide and 5-Year Mortality Risk in Stable Coronary Artery Disease: The Contributory Role of Intestinal Microbiota in a COURAGE-Like Patient Cohort. J Am Heart Assoc 5, doi:10.1161/JAHA.115.002816 (2016).

12 Ross, F. C. et al. The interplay between diet and the gut microbiome: implications for health and disease. Nat Rev Microbiol 22, 671–686, doi:10.1038/s41579-024-01068-4 (2024).

13 Key, T. J., Papier, K. & Tong, T. Y. N. Plant-based diets and long-term health: findings from the EPIC-Oxford study. Proc Nutr Soc 81, 190–198, doi:10.1017/S0029665121003748 (2022).

14 Yannakoulia, M. & Scarmeas, N. Diets. N Engl J Med 390, 2098–2106, doi:10.1056/NEJMra2211889 (2024).

15 Rouskas, K. et al. Periodic dietary restriction of animal products induces metabolic reprogramming in humans with effects on cardiometabolic health. npj Metabolic Health and Disease 3, 14, doi:10.1038/s44324-025-00057-2 (2025).

16 Loizidou, E. M. et al. Short-term animal product restriction alters metabolic profiles and modulates immune function. medRxiv, 2025.2005.2023.25328246, doi:10.1101/2025.05.23.25328246 (2025).

17 Simistiras, A. et al. Diet-responsive proteogenomic effects following short-term restriction of animal products in humans. bioRxiv, 2025.2007.2021.665884, doi:10.1101/2025.07.21.665884 (2025).

18 Georgakouli, K. et al. The Effects of Greek Orthodox Christian Fasting during Holy Week on Body Composition and Cardiometabolic Parameters in Overweight Adults. Diseases 10, doi:10.3390/diseases10040120 (2022).

19 Sarri, K. O., Linardakis, M. K., Bervanaki, F. N., Tzanakis, N. E. & Kafatos, A. G. Greek Orthodox fasting rituals: a hidden characteristic of the Mediterranean diet of Crete. Br J Nutr 92, 277–284, doi:10.1079/BJN20041197 (2004).

20 R Core Team. R: A Language and Environment for Statistical Computing. R Foundation for Statistical Computing (2023).

21 Andrews, S. FastQC: a quality control tool for high throughput sequence data, <http://www.bioinformatics.babraham.ac.uk/projects/fastqc> (2010).

22 Ewels, P., Magnusson, M., Lundin, S. & Kaller, M. MultiQC: summarize analysis results for multiple tools and samples in a single report. Bioinformatics 32, 3047–3048, doi:10.1093/bioinformatics/btw354 (2016).

23 Martin, M. Cutadapt removes adapter sequences from high-throughput sequencing reads. EMBnet.journal 17, 10–12, 10.14806/ej.17.1.200. (2011).

24 Li, H. Aligning sequence reads, clone sequences and assembly contigs with BWA-MEM. arXiv, 10.48550/arXiv.1303.3997 (2013).

25 Callahan, B. J. et al. DADA2: High-resolution sample inference from Illumina amplicon data. Nat Methods 13, 581–583, doi:10.1038/nmeth.3869 (2016).

26 Quast, C. et al. The SILVA ribosomal RNA gene database project: improved data processing and web-based tools. Nucleic Acids Res 41, D590–596, doi:10.1093/nar/gks1219 (2013).

27 McLaren, M. R. & Callahan, B. J. Silva 138.1 prokaryotic SSU taxonomic training data formatted for DADA2 [Data set]. Zenodo, Geneva, Switzerland (2021).

28 McMurdie, P. J. & Holmes, S. phyloseq: an R package for reproducible interactive analysis and graphics of microbiome census data. PLoS One 8, e61217, doi:10.1371/journal.pone.0061217 (2013).

29 Oksanen, J., Simpson, G., Blanchet, F., Kindt, R., Legendre, P., Minchin, P., O’Hara, R., Solymos, P., Stevens, M., Szoecs, E., Wagner, H., Barbour, M., Bedward, M., Bolker, B., Borcard, D., Carvalho, G., Chirico, M., De Caceres, M., Durand, S., Evangelista, H., FitzJohn, R., Friendly, M., Furneaux, B., Hannigan, G., Hill, M., Lahti, L., McGlinn, D., Ouellette, M., Ribeiro, Cunha, E., Smith, T., Stier, A., Ter Braak, C., Weedon, J. vegan: Community Ecology Package. R package version 2.6-8, doi:https://CRAN.R-project.org/package=vegan (2024).

30 Anderson, M. J. A new method for non-parametric multivariate analysis of variance. Austral Ecology 26, 32–46, 10.1111/j.1442-9993.2001.01070.pp.x (2001).

31 Mallick, H. et al. Multivariable association discovery in population-scale meta-omics studies. PLOS Computational Biology 17, e1009442, doi:10.1371/journal.pcbi.1009442 (2021).

32 Langille, M. G. et al. Predictive functional profiling of microbial communities using 16S rRNA marker gene sequences. Nat Biotechnol 31, 814–821, doi:10.1038/nbt.2676 (2013).

33 Singh, A. et al. DIABLO: an integrative approach for identifying key molecular drivers from multi-omics assays. Bioinformatics 35, 3055–3062, doi:10.1093/bioinformatics/bty1054 (2019).

34 Rohart, F., Gautier, B., Singh, A. & Le Cao, K. A. mixOmics: An R package for ’omics feature selection and multiple data integration. PLoS Comput Biol 13, e1005752, doi:10.1371/journal.pcbi.1005752 (2017).

35 Rezac, S., Kok, C. R., Heermann, M. & Hutkins, R. Fermented Foods as a Dietary Source of Live Organisms. Frontiers in microbiology 9, 1785, doi:10.3389/fmicb.2018.01785 (2018).

36 Lee, C. et al. Effect of Consumption of Animal Products on the Gut Microbiome Composition and Gut Health. Food Sci Anim Resour 43, 723–750, doi:10.5851/kosfa.2023.e44 (2023).

37 Palego, L., Betti, L., Rossi, A. & Giannaccini, G. Tryptophan Biochemistry: Structural, Nutritional, Metabolic, and Medical Aspects in Humans. J Amino Acids 2016, 8952520, doi:10.1155/2016/8952520 (2016).

38 Thakur, K., Tomar, S. K., Singh, A. K., Mandal, S. & Arora, S. Riboflavin and health: A review of recent human research. Crit Rev Food Sci Nutr 57, 3650–3660, doi:10.1080/10408398.2016.1145104 (2017).

39 TeSlaa, T., Ralser, M., Fan, J. & Rabinowitz, J. D. The pentose phosphate pathway in health and disease. Nat Metab 5, 1275–1289, doi:10.1038/s42255-023-00863-2 (2023).

40 Al-Refai, W., Keenan, S., Camera, D. M. & Cooke, M. B. The Influence of Vegan, Vegetarian, and Omnivorous Diets on Protein Metabolism: A Role for the Gut-Muscle Axis? Nutrients 17, doi:10.3390/nu17071142 (2025).

41 Seel, W., Reiners, S., Kipp, K., Simon, M.-C. & Dawczynski, C. Role of Dietary Fiber and Energy Intake on Gut Microbiome in Vegans, Vegetarians, and Flexitarians in Comparison to Omnivores—Insights from the Nutritional Evaluation (NuEva) Study. Nutrients 15, 1914 (2023).

42 Prochazkova, M. et al. Vegan Diet Is Associated With Favorable Effects on the Metabolic Performance of Intestinal Microbiota: A Cross-Sectional Multi-Omics Study. Frontiers in Nutrition 8, doi:10.3389/fnut.2021.783302 (2022).

43 van der Vossen, E. W. J. et al. Gut microbiome transitions across generations in different ethnicities in an urban setting-the HELIUS study. Microbiome 11, 99, doi:10.1186/s40168-023-01488-z (2023).

44 Jousset, A. et al. Where less may be more: how the rare biosphere pulls ecosystems strings. ISME J 11, 853–862, doi:10.1038/ismej.2016.174 (2017).

45 Latorre-Pérez, A. et al. The Spanish gut microbiome reveals links between microorganisms and Mediterranean diet. Scientific Reports 11, 21602, doi:10.1038/s41598-021-01002-1 (2021).

46 van Soest, A. P. M. et al. Associations between Pro- and Anti-Inflammatory Gastro-Intestinal Microbiota, Diet, and Cognitive Functioning in Dutch Healthy Older Adults: The NU-AGE Study. Nutrients 12, doi:10.3390/nu12113471 (2020).

47 De Angelis, M. et al. Diet influences the functions of the human intestinal microbiome. Scientific Reports 10, 4247, doi:10.1038/s41598-020-61192-y (2020).

48 Asnicar, F. et al. Microbiome connections with host metabolism and habitual diet from 1,098 deeply phenotyped individuals. Nat Med 27, 321–332, doi:10.1038/s41591-020-01183-8 (2021).

49 Wang, S. et al. Microbial collaborations and conflicts: unraveling interactions in the gut ecosystem. Gut Microbes 16, 2296603, doi:10.1080/19490976.2023.2296603 (2024).

50 Losno, E. A., Sieferle, K., Perez-Cueto, F. J. A. & Ritz, C. Vegan Diet and the Gut Microbiota Composition in Healthy Adults. Nutrients 13, 2402 (2021).

51 van Faassen, A. et al. Bile acids, neutral steroids, and bacteria in feces as affected by a mixed, a lacto-ovovegetarian, and a vegan diet. Am J Clin Nutr 46, 962–967, doi:10.1093/ajcn/46.6.962 (1987).

52 Chen, Y. et al. Exploiting lactic acid bacteria for inflammatory bowel disease: A recent update. Trends in Food Science & Technology 138, 126–140, 10.1016/j.tifs.2023.06.007 (2023).

53 De Filippis, F., Pasolli, E. & Ercolini, D. The food-gut axis: lactic acid bacteria and their link to food, the gut microbiome and human health. FEMS Microbiol Rev 44, 454–489, doi:10.1093/femsre/fuaa015 (2020).

54 Gryaznova, M. V. et al. Study of microbiome changes in patients with ulcerative colitis in the Central European part of Russia. Heliyon 7, e06432, doi:10.1016/j.heliyon.2021.e06432 (2021).

55 Mai, H. et al. The role of gut microbiota in the occurrence and progression of non-alcoholic fatty liver disease. Frontiers in microbiology 14, 1257903, doi:10.3389/fmicb.2023.1257903 (2023).

56 Dovrolis, N. et al. The Interplay between Mucosal Microbiota Composition and Host Gene-Expression is Linked with Infliximab Response in Inflammatory Bowel Diseases. Microorganisms 8, doi:10.3390/microorganisms8030438 (2020).

57 Wu, I. W. et al. Integrative metagenomic and metabolomic analyses reveal severity-specific signatures of gut microbiota in chronic kidney disease. Theranostics 10, 5398–5411, doi:10.7150/thno.41725 (2020).

58 Dahal, R. H., Kim, S., Kim, Y. K., Kim, E. S. & Kim, J. Insight into gut dysbiosis of patients with inflammatory bowel disease and ischemic colitis. Frontiers in microbiology 14, 1174832, doi:10.3389/fmicb.2023.1174832 (2023).

59 Molinas-Vera, M., Ferreira-Sanabria, G., Peña, P., Sandoval-Espinola, W.J. The Paraguayan gut microbiome contains high abundance of the phylumActinobacteriota and reveals the influence of health and lifestyle factors. Gut microbes report 1, doi:0.1080/29933935.2024.2332988 (2024).

60 Guo, G. et al. Exploring the causal effects of the gut microbiome on serum lipid levels: A two-sample Mendelian randomization analysis. Frontiers in microbiology 14, 1113334, doi:10.3389/fmicb.2023.1113334 (2023).

61 Sheng, S. et al. Gut microbiome is associated with metabolic syndrome accompanied by elevated gamma-glutamyl transpeptidase in men. Front Cell Infect Microbiol 12, 946757, doi:10.3389/fcimb.2022.946757 (2022).

62 Lun, H. et al. Altered gut microbiota and microbial biomarkers associated with chronic kidney disease. Microbiologyopen 8, e00678, doi:10.1002/mbo3.678 (2019).

63 Van Hul, M. et al. What defines a healthy gut microbiome? Gut 73, 1893–1908, doi:10.1136/gutjnl-2024-333378 (2024).

64 Guerra, B. et al. The Mevalonate Pathway, a Metabolic Target in Cancer Therapy. Front Oncol 11, 626971, doi:10.3389/fonc.2021.626971 (2021).

65 Luo, J., Yang, H. & Song, B. L. Mechanisms and regulation of cholesterol homeostasis. Nat Rev Mol Cell Biol 21, 225–245, doi:10.1038/s41580-019-0190-7 (2020).

66 Buhaescu, I. & Izzedine, H. Mevalonate pathway: a review of clinical and therapeutical implications. Clin Biochem 40, 575–584, doi:10.1016/j.clinbiochem.2007.03.016 (2007).

67 Wang, C. H., Hou, J., Deng, H. K. & Wang, L. J. Microbial production of mevalonate. J Biotechnol 370, 1–11, doi:10.1016/j.jbiotec.2023.05.005 (2023).

68 Weiner, I. D., Mitch, W. E. & Sands, J. M. Urea and Ammonia Metabolism and the Control of Renal Nitrogen Excretion. Clin J Am Soc Nephrol 10, 1444–1458, doi:10.2215/CJN.10311013 (2015).

69 Valles, C. et al. Negativibacillus massiliensis gen. nov., sp. nov., a New Bacterial Genus Isolated from a Human Left Colon Sample. Microbiology Research 12, 29–42 (2021).

70 Ginsberg, H. N. et al. Triglyceride-rich lipoproteins and their remnants: metabolic insights, role in atherosclerotic cardiovascular disease, and emerging therapeutic strategies-a consensus statement from the European Atherosclerosis Society. Eur Heart J 42, 4791–4806, doi:10.1093/eurheartj/ehab551 (2021).

71 Yang, X. H., Zhang, B. L., Cheng, Y., Fu, S. K. & Jin, H. M. Association of remnant cholesterol with risk of cardiovascular disease events, stroke, and mortality: A systemic review and meta-analysis. Atherosclerosis 371, 21–31, doi:10.1016/j.atherosclerosis.2023.03.012 (2023).

72 Flippo, K. H. & Potthoff, M. J. Metabolic Messengers: FGF21. Nat Metab 3, 309–317, doi:10.1038/s42255-021-00354-2 (2021).

73 Martin, A. et al. Gut microbiota mediate the FGF21 adaptive stress response to chronic dietary protein-restriction in mice. Nat Commun 12, 3838, doi:10.1038/s41467-021-24074-z (2021).

74 Liu, S. et al. A Comprehensive Analysis of HAVCR1 as a Prognostic and Diagnostic Marker for Pan-Cancer. Front Genet 13, 904114, doi:10.3389/fgene.2022.904114 (2022).

75 Zhang, J. et al. The biological functions and related signaling pathways of SPON2. Front Oncol 13, 1323744, doi:10.3389/fonc.2023.1323744 (2023).

76 Zhu, L. H. et al. Mindin/Spondin 2 inhibits hepatic steatosis, insulin resistance, and obesity via interaction with peroxisome proliferator-activated receptor alpha in mice. J Hepatol 60, 1046–1054, doi:10.1016/j.jhep.2014.01.011 (2014).

77 Sorci, L. et al. Initial-Rate Kinetics of Human NMN-Adenylyltransferases:[] Substrate and Metal Ion Specificity, Inhibition by Products and Multisubstrate Analogues, and Isozyme Contributions to NAD+ Biosynthesis. Biochemistry 46, 4912–4922, doi:10.1021/bi6023379 (2007).

78 Georgiadis, M. M. et al. Evolution of the redox function in mammalian apurinic/apyrimidinic endonuclease. Mutation Research/Fundamental and Molecular Mechanisms of Mutagenesis 643, 54–63, 10.1016/j.mrfmmm.2008.04.008 (2008).

79 Schilperoort, M., Ngai, D., Katerelos, M., Power, D. A. & Tabas, I. PFKFB2-mediated glycolysis promotes lactate-driven continual efferocytosis by macrophages. Nat Metab 5, 431–444, doi:10.1038/s42255-023-00736-8 (2023).

80 Wu, H. et al. Syndecan-4 shedding is involved in the oxidative stress and inflammatory responses in left atrial tissue with valvular atrial fibrillation. Int J Clin Exp Pathol 8, 6387–6396 (2015).

81 Kelly, T. N. et al. Gut Microbiome Associates With Lifetime Cardiovascular Disease Risk Profile Among Bogalusa Heart Study Participants. Circ Res 119, 956–964, doi:10.1161/CIRCRESAHA.116.309219 (2016).

82 Olaisen, M. et al. Bacterial Mucosa-associated Microbiome in Inflamed and Proximal Noninflamed Ileum of Patients With Crohn’s Disease. Inflamm Bowel Dis 27, 12–24, doi:10.1093/ibd/izaa107 (2021).

83 Truong, T. Q., Falstrault, L., Tremblay, C. & Brissette, L. Low density lipoprotein-receptor plays a major role in the binding of very low density lipoproteins and their remnants on HepG2 cells. Int J Biochem Cell Biol 31, 695–705, doi:10.1016/s1357-2725(99)00014-x (1999).

84 Jeong, M. H. et al. Cdon deficiency causes cardiac remodeling through hyperactivation of WNT/beta-catenin signaling. Proc Natl Acad Sci U S A 114, E1345–E1354, doi:10.1073/pnas.1615105114 (2017).

85 Vo, T. T. et al. Exploring scavenger receptor class F member 2 and the importance of scavenger receptor family in prediagnostic diseases. Toxicol Res 39, 341–353, doi:10.1007/s43188-023-00176-2 (2023).

86 Lin, X. et al. Integrin beta5 subunit regulates hyperglycemia-induced vascular endothelial cell apoptosis through FoxO1-mediated macroautophagy. Chin Med J (Engl) 137, 565–576, doi:10.1097/CM9.0000000000002769 (2024).

87 Tang, J. et al. The therapeutic value of bifidobacteria in cardiovascular disease. NPJ Biofilms Microbiomes 9, 82, doi:10.1038/s41522-023-00448-7 (2023).

88 Wang, Z. et al. Gut flora metabolism of phosphatidylcholine promotes cardiovascular disease. Nature 472, 57–63, doi:10.1038/nature09922 (2011).

89 Iram, T. et al. Megf10 Is a Receptor for C1Q That Mediates Clearance of Apoptotic Cells by Astrocytes. J Neurosci 36, 5185–5192, doi:10.1523/JNEUROSCI.3850-15.2016 (2016).

90 Carmody, R. N., Varady, K. & Turnbaugh, P. J. Digesting the complex metabolic effects of diet on the host and microbiome. Cell 187, 3857–3876, doi:10.1016/j.cell.2024.06.032 (2024).

