## Supplementary Material for "Rapid remodeling of the human gut microbiome in response to short-term animal product restriction"

**Supplementary Text 1.** **FastBio study inclusion and exclusion criteria.**

Inclusion criteria:

- Healthy female and male subjects 18 – 75 years of age at the time of enrolment. One participant turned 76 between time of enrolment and first sampling timepoint (T1).
- Able to provide signed and dated informed consent. Willing to provide blood samples.
- Individuals who had practiced periodic animal product restriction for at least ten years (for the PR group).
- Individuals who had not practiced any kind of specific diet including veganism, vegetar-ianism, caloric restriction, intermittent fasting (for the NR group).

Exclusion criteria:

- Use of antibiotic, antifungal, antiviral or antiparasitic drugs six months prior to T1.
- Acute disease, defined as the presence of a moderate or severe illness with or without fever, at sampling timepoints.
- Alcohol or drug abuse two years prior to T1.
- Participants who are normally periodically abstaining from animal products, but had to alter their diet for a specific reason (e.g. pregnancy).

**Supplementary Text 2. Selected covariates.**

For the differential abundance analyses in MaAsLin2, we added sex, age^2^, BMI, medication use, smoking and Bristol score as fixed effects in the LM and CPLM models.

For the correlation analyses between bacterial abundance and levels of blood biomarkers and molecular phenotypes, we used linear regression modeling to regress out the effects of:

(i) age^2^, sex, BMI, medication use, smoking and Bristol score from bacterial abundance
(ii) age^2^, sex, BMI, medication use and smoking from biomarker levels
(iii) age^2^, sex, BMI, medication use and smoking from metabolite levels
(iv) age^2^, sex, BMI, medication use, smoking and mean protein value from protein levels

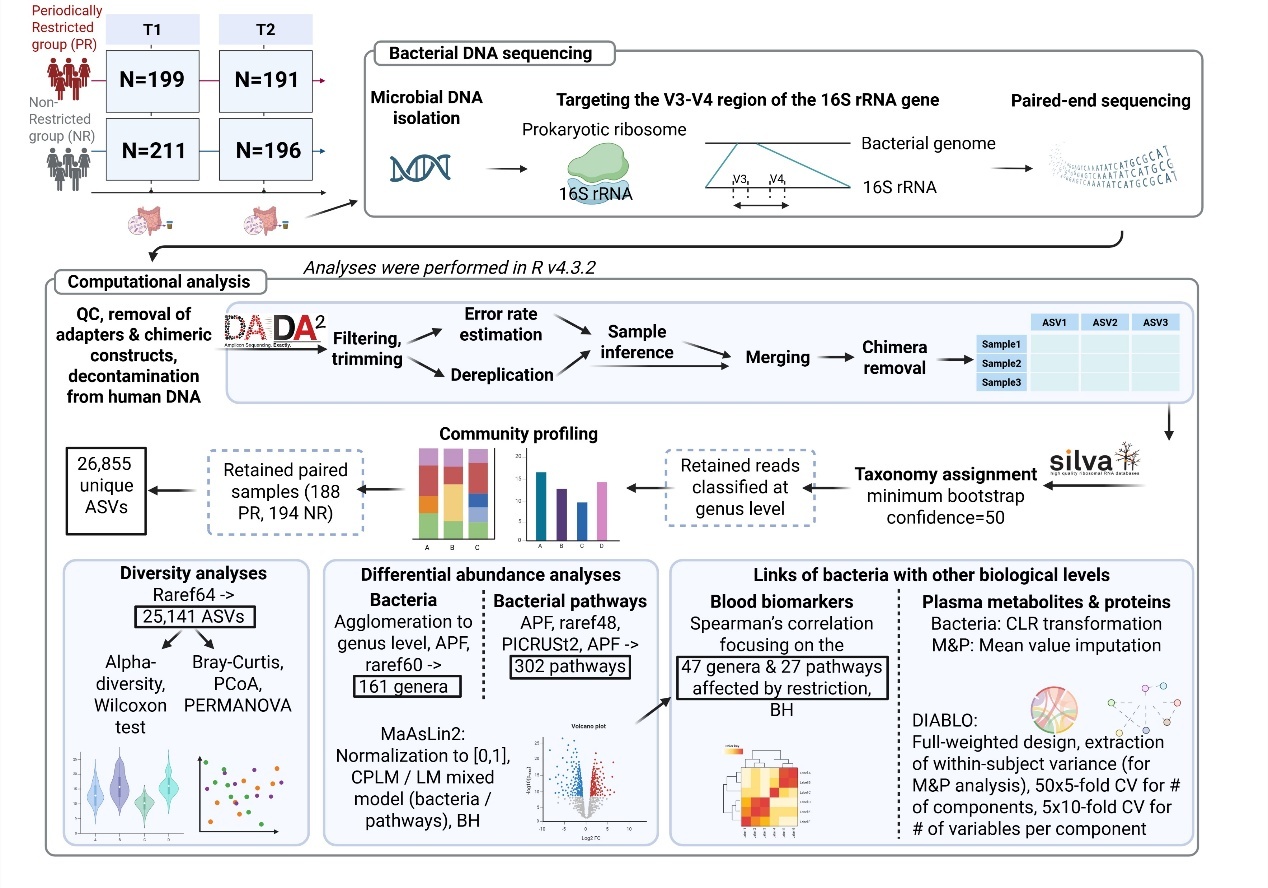

**Supplementary Figure 1. Analysis workflow.** APF: abundance (0.0001%) and prevalence (10%) filtering; RarefN: Rarefaction to N thousand reads; BH: Benjamini-Hochberg method for multiple testing correction; M&P: Metabolites and proteins. All significance thresholds are equal to 0.05. Created using BioRender.com.

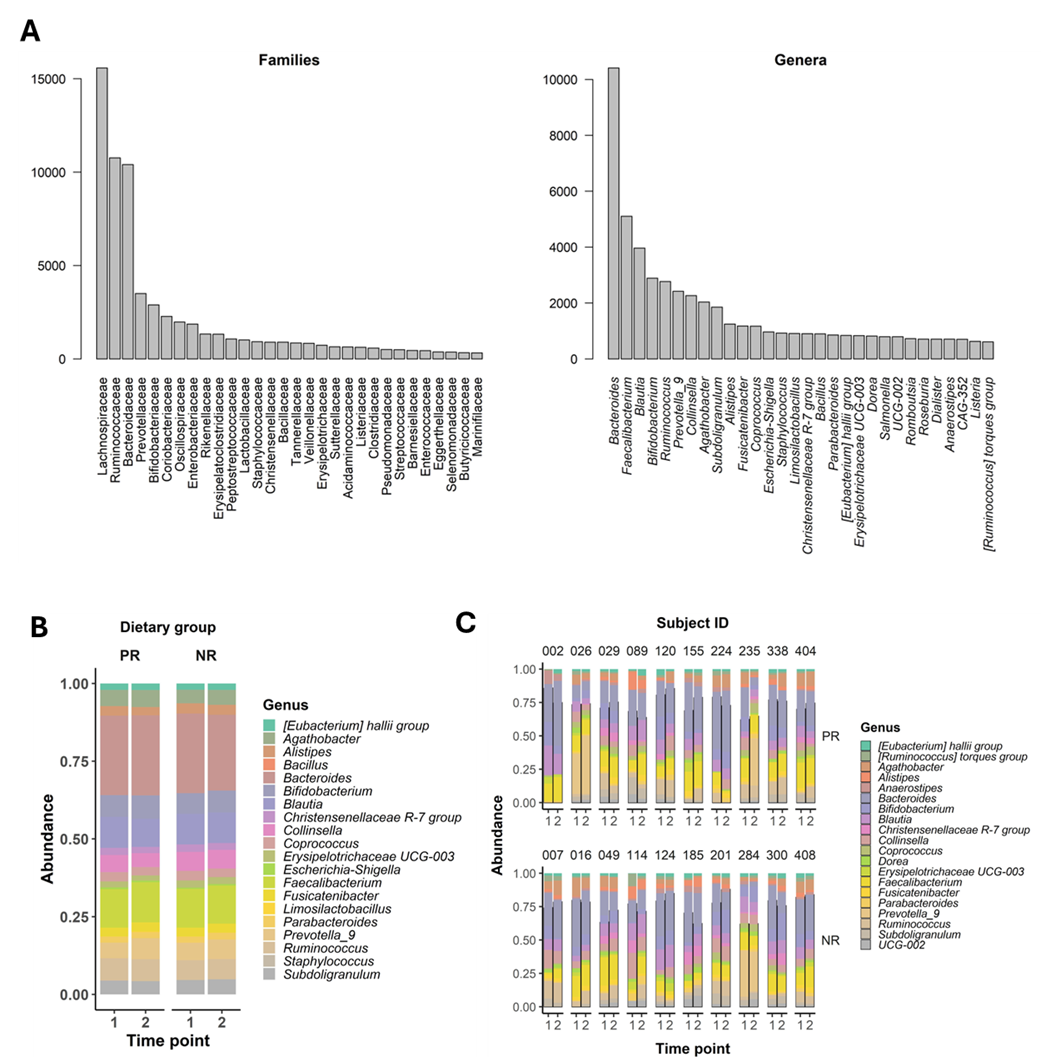

**Supplementary Figure 2. Taxonomic profiling. A**. Bar plots showing the average abundance per sample of the 30 most dominant families and genera. **B, C**. Stacked bar plots showing the relative abundance of the 20 most abundant genera, in all individuals grouped by dietary group and time point (**B**) and in ten participants from each dietary group during each time point (**C**).

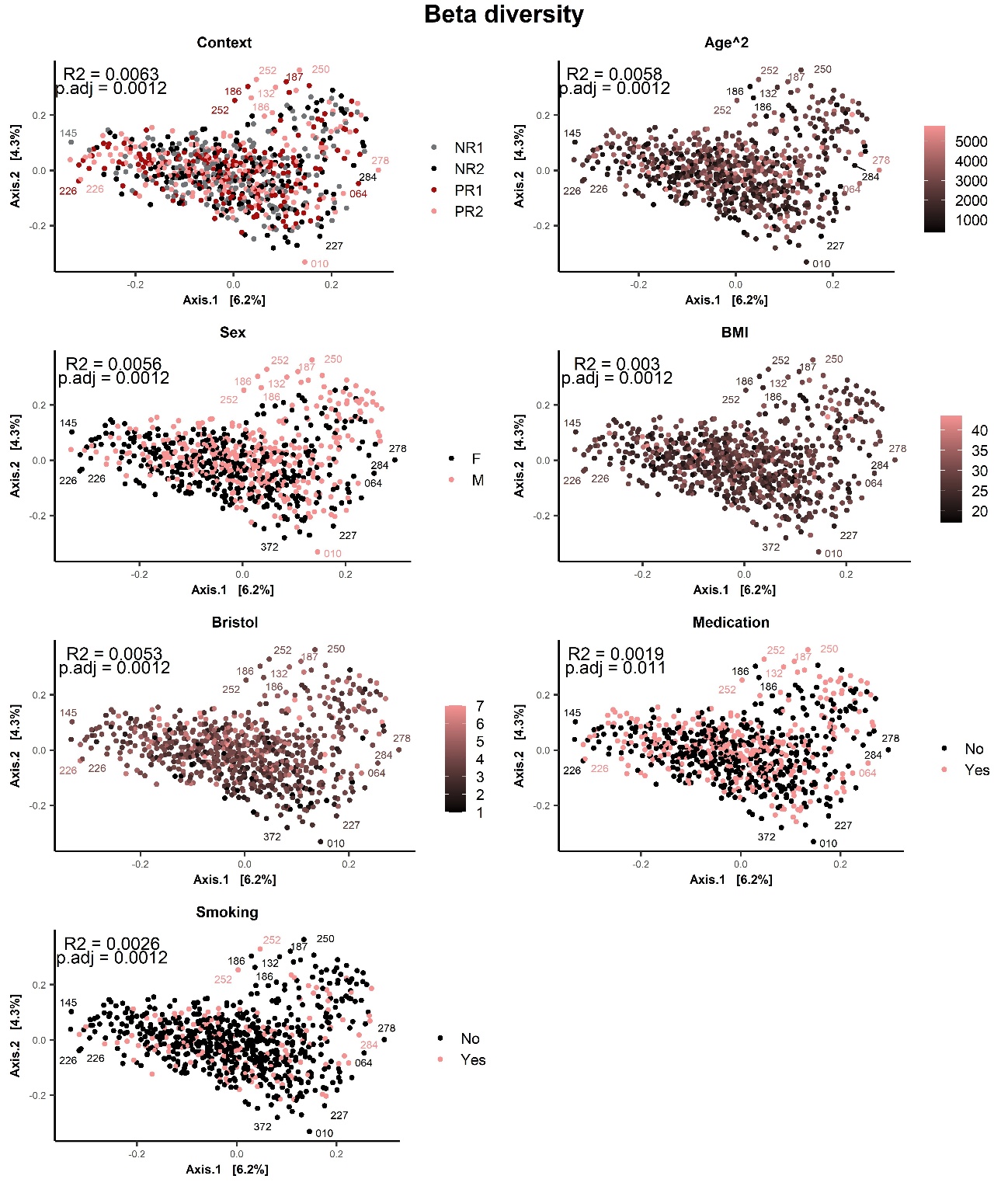

**Supplementary Figure 3. Beta diversity.** Principal Coordinates Analysis (PCoA) based on Bray-Curtis dissimilarity, showing differences in bacterial composition between samples. Each dot represents one sample and is colored according to the value of each explanatory variable. Statistical significance of group separation was assessed with PERMANOVA using a sequential model. The coefficients of determination (R2) and p-values (BH-adjusted) for 999 permutations are shown.

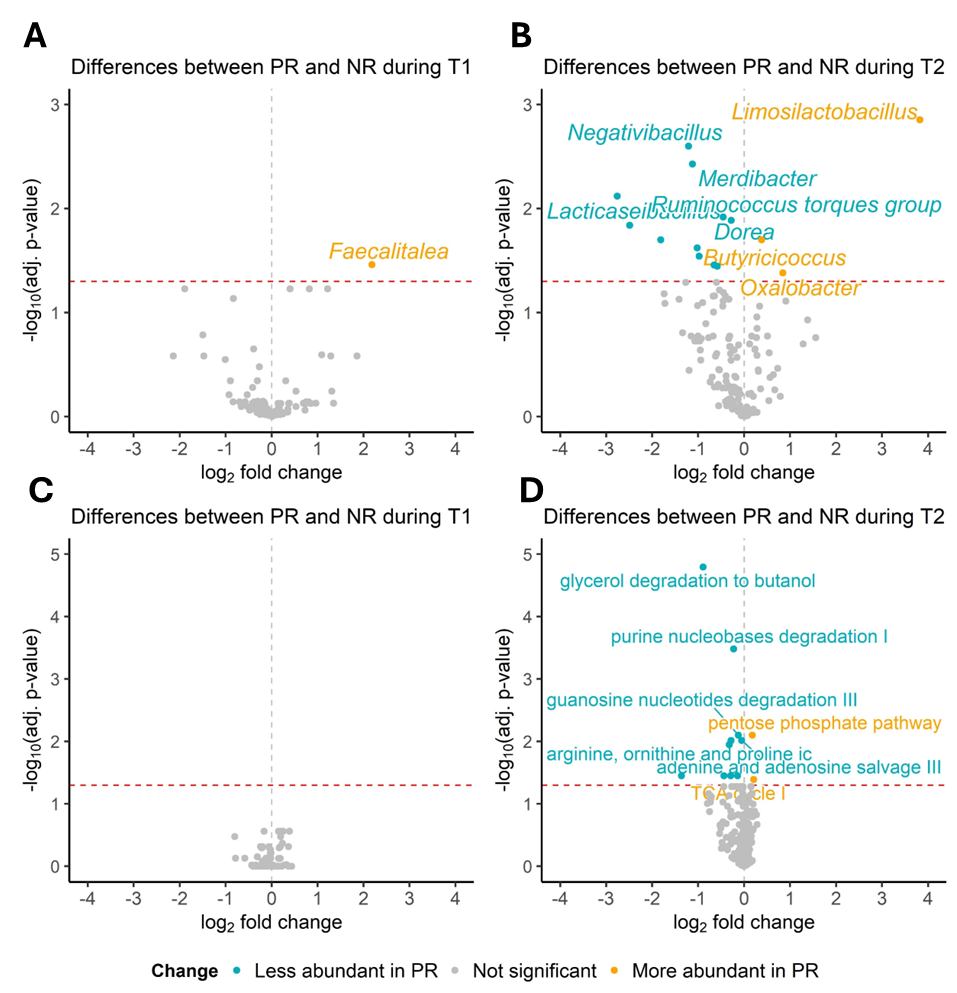

**Supplementary Figure 4. Differentially abundant genera and pathways between dietary groups.** Volcano plots displaying log2 fold changes and -log10 p-values (BH-adjusted) for bacterial genera between dietary groups at time point 1 (**A**) and at time point 2 (**B**), and for microbially expressed predicted pathways between dietary groups at time point 1 (**C**) and at time point 2 (**D**), Significant changes (p<0.05) are highlighted in blue (less abundant in PR) or yellow (more abundant in PR) color. In each direction, up to five most significant genera and pathways are labeled. Ic: interconversion.

|  | **Total (N=411)** | **PR (N=200)** | **NR (N=211)** | ***P*-value** |
| --- | --- | --- | --- | --- |
| **Sex** |  |  |  |  |
| Female | 224 (54.5) | 108 (54) | 116 (55) | 0.8425 |
| Male | 187 (45.5) | 92 (46) | 95 (45) |  |
| **Age (yrs)** | 48.1 ± 13.6 | 51.5 ± 13.5 | 45.0 ± 13.1 | <0.00001 |
| **BMI (Kg/m^2^)** | 27.3 ± 4.6 | 28.4 ± 4.6 | 26.2 ± 4.4 | <0.00001 |
| **Blood Pressure (mmHg)** |  |  |  |  |
| Systolic BP (SBP) | 124 ± 19.4 | 127 ± 19.0 | 121 ± 19.7 | 0.003 |
| Diastolic BP (DBP) | 79 ± 11.3 | 80 ± 10.8 | 78 ± 11.8 | 0.04 |
| **Education** |  |  |  |  |
| Tertiary | 293 (71.3) | 134 (67) | 159 (75.4) | 0.0612 |
| Primary and Secondary | 118 (28.7) | 66 (33) | 52 (24.6) |  |
| **Marital status** |  |  |  |  |
| Married | 286 (69.6) | 147 (73.5) | 139 (65.9) | 0.0931 |
| Unmarried | 125 (30.4) | 53 (26.5) | 72 (34.1) |  |
| **Smoking (Y/N)** |  |  |  |  |
| Non-smokers | 328 (79.8) | 187 (93.5) | 141 (66.8) | <0.00001 |
| Smokers | 83 (20.2) | 13 (6.5) | 70 (33.2) |  |
| **Parental origin** |  |  |  |  |
| Northern Greece | 275 (66.9) | 137 (68.5) | 138 (65.4) | 0.0979 |
| Central or Southern Greece | 32 (7.8) | 11 (5.5) | 21 (9.9) |  |

**Supplementary Table 1. Sociodemographic traits and geographic origin of FastBio study participants.** The FastBio population sample has been described in detail in (Rouskas et al., 2025). Continuous variables were expressed as mean ± standard deviations and categorical variables as N (%). SBP = Systolic Blood Pressure; DBP = Diastolic Blood Pressure. P-values are from chi-square test or Mann-Whitney test for categorical and numerical variables, respectively. Stool samples were delivered by 199 PR and 211 NR individuals.

**Supplementary Table 2**. **Differentially abundant genera and pathways across contexts.** (xlsx)

| **Family** | **Genus** | **Direction** |
| --- | --- | --- |
| Anaerovoracaceae | *Eubacterium nodatum group* |  |
|  | *Family XIII AD3011 group* |  |
| Butyricicoccaceae | *Butyricicoccus* |  |
|  | *UCG-008* |  |
| Coriobacteriaceae | *Enorma* |  |
|  | *Collinsella* |  |
| Erysipelatoclostridiaceae | *UCG-004* |  |
|  | *Erysipelatoclostridium* |  |
| Erysipelotrichaceae | *Holdemanella* |  |
|  | *Holdemania* |  |
|  | *Clostridium innocuum group* |  |
|  | *Merdibacter* |  |
|  | *Faecalitalea* |  |
|  | *Turicibacter* |  |
| Lachnospiraceae | *Ruminococcus torques group* |  |
|  | *Dorea* |  |
|  | *Ruminococcus gnavus group* |  |
|  | *Lachnospira* |  |
|  | *AC2044 group* |  |
|  | *Tyzzerella* |  |
|  | *UCG-010* |  |
|  | *Frisingicoccus* |  |
|  | *Blautia* |  |
| Lactobacillaceae | *Lactiplantibacillus* |  |
|  | *Lacticaseibacillus* |  |
|  | *Lactobacillus* |  |
|  | *Latilactobacillus* |  |
|  | *Ligilactobacillus* |  |
| Oscillospiraceae | *NK4A214 group* |  |
|  | *Intestinimonas* |  |
|  | *Hydrogenoanaerobacterium* |  |
|  | *Flavonifractor* |  |
| Peptostreptococcaceae | *Terrisporobacter* |  |
|  | *Intestinibacter* |  |
| Ruminococcaceae | *Negativibacillus* |  |
|  | *UBA1819* |  |
|  | *Pygmaiobacter* |  |
| Streptococcaceae | *Streptococcus* |  |
|  | *Lactococcus* |  |

**Supplementary Table 3. Direction of changes in abundance upon dietary restriction for members of families that encompass more than one affected member.**

**References**

Rouskas, K., Bocher, O., Simistiras, A., Emmanouil, C., Mantas, P., Skoulakis, A., Park, Y.-C., Dimopoulos, A., Glentis, S., Kastenmüller, G.*, et al.* (2025). Periodic dietary restriction of animal products induces metabolic reprogramming in humans with effects on cardiometabolic health. npj Metabolic Health and Disease *3*, 14.
